# Divergent epigenetic profiles from two differentially impacted wild populations of estuarine cordgrass (*Sporobolus alterniflorus*)

**DOI:** 10.1101/2021.03.22.436412

**Authors:** L. DeCarlo, F. Meckler, M. Hans, S. Kelemen, H. Magun, M. Noah, L. Pappajohn, N. Anderson, R. Berger, J. Berkel, N. Brooke, L. Chen, O. Chijioke, N. Dewees, P. Falkner, J. Frank, W. Holzman, V. Marino, A. Ravaschiere, Y. Wang, A. Williams, Z. Williams, D. Gentile, R.L. Cox

## Abstract

The effects of urbanization on watershed ecosystems present critical challenges to modern survival. Organisms in urbanized areas experience high rates of evolutionary change, but genetic adaptation alone cannot mitigate the rapid and severe effects of urbanization on biodiversity. Highly resilient, foundation species are key to maintaining an ecosystem’s integrity in the face of urban stressors. However, the rapid collapse and disappearance of watershed ecosystems calls into question the extent to which we can rely on such species for their services. Our research investigates the molecular mechanisms by which the foundation ecosystems provider, *Sporobolus alterniflorus*, adapts to life in an urbanized environment. To elucidate these mechanisms, we quantified changes in global DNA methylation (% 5-mC) as a result of acute heat stress. Specimens from two differentially impacted populations across an urban to suburban geographical transect formed the basis of this study. These two populations of *Sporobolus alterniflora* exhibit inverse global DNA methylation patterns when exposed to the same acute heat stress. Our findings suggest that epigenetic mechanisms, such as DNA methylation, control rapid and transient adaptation, in the form of differential stress responses, to distinct environment challenges.

**Highlights for manuscript submission:**

▪ estuarine grasses native to the Bronx River, NY face stresses associated with low dissolved oxygen and urbanization
▪ differentially impacted populations of estuarine grasses exhibit inverse global DNA methylation profiles in response to acute heat stress
▪ DNA methylation may represent a mechanism by which plants transiently respond to environmental stressors, and this may represent a form of rapid adaptive evolution
▪ stress priming by transgenerational epigenetic modification may enhance fitness in grasses native to the heavily impacted Bronx River estuary

## 1. Introduction

Urban estuaries are highly productive ecosystems that provide critical services including storm surge protection, water filtration and wildlife habitat (Barbier et al., 2011; Franca et al., 2012). Coastal salt marshes comprise approximately 25% of the global soil carbon sink through plant production and high carbon burial rates (Chmura et al., 2003). Human activity, industrialization, and climate change continue to negatively impact estuarine ecosystems, especially in urbanized areas (Limburg et al., 2005; Astaraie-Imani et al., 2012; Chin et al., 2013). Estuarine species face stresses associated with adverse environmental conditions such as low dissolved oxygen, combined sewage overflow, toxin contamination, bank destabilization, habitat degradation, and extreme temperature fluctuation (Van Dolah, et al., 2008; Courrat et al., 2009; Halem et al., 2014; Ravaschiere et al., 2017). Knowledge of the molecular mechanisms underlying stress adaptation is essential if we hope to rehabilitate estuarine ecosystems adversely affected by urbanization.

Urbanization is often associated with rapidly changing anthropogenic stressors (Alberti, 2015; Donihue and Lambert, 2015). As sessile organisms, plants must respond to these environmental challenges by rapidly regulating gene expression, and this is often accomplished via epigenetic alterations such as DNA methylation and histone modification (Arikan et al., 2018; Wang et al., 2010). These transient and rapid modifications provide essential “on demand” phenotypic variation (Rey et al., 2016), that may represent critical adaptive mechanisms for species such as marsh grasses that provide essential ecological services to urban communities.

Here, we take an *in vivo* approach to the study of epigenetic adaptation in wild populations of smooth cordgrass *Sporobolus alterniflorus* (formerly *Spartina alterniflora*), a critical ecosystems provider and foundation species that is native to estuaries throughout the North American east coast (Gedan and Bertness, 2010; Peterson, 2014). This halophyte species is particularly resilient in the face of urbanization (Gedan and Bertness, 2010) and thus represents a particularly suitable model for our study. In research findings presented here, we track two robust mechanisms for transient stress response in plants: genomic DNA cytosine methylation (% 5-mC) and heat shock protein 70 expression (HSP70). The methylation of 5-cytosine (% 5-mC) occurs when a methyl group is enzymatically attached to the 5’ carbon of cytosine’s pyrimidine ring, resulting in a decrease in gene expression at the methylation site. This reversible chemical modification elicits transient phenotypic changes required for stress response, especially in plants (Meyer, 2008; Boyko and Kovalchuk, 2011; Arikan et al., 2018). HSP70 chaperone protein is a well-characterized and universal stress responder that facilitates refolding of denatured proteins, and thus restoration of vital metabolic activities (Walter and Ron, 2011).

We compared global DNA methylation (% 5-mC) in response to acute heat stress in two differentially impacted wild populations: the Bronx River estuary, New York and the Greenwich Cove estuary, Connecticut, using HSP70 as an indicator of acute stress response. These estuaries are located about 25 miles from each other and share similar biota. However, their diametric histories make these estuaries uniquely suited for a comparative analysis of urban stress response. The Bronx River estuary has a history of factory waste and sewage dumping (Crimmens and Larson, 2006). Construction of the Bronx River Parkway led to degradation of water quality and a decrease in species diversity (Rachlin, 2007). Ongoing industrialization eventually led to benzo-a pyrene (Litten et al., 2007) and sewage contamination (Rachlin, 2007; Wang and Pant, 2010) of the Bronx River estuary. Whereas, the suburban Greenwich Cove estuary lies at a greater geographic distance from urbanized centers and thus experiences comparatively less impact from urbanization and industrialization (Halem et al., 2014; Ravaschiere et al., 2017). Previous studies of the Bronx River estuary document endocrine disruption (Halem et al., 2014) and altered heat shock response (Ravaschiere et al., 2017) in the native ecosystem provider (Galimay et. al., 2017) Atlantic ribbed mussel (*Geukensia demissa*). In addition, our nine-year longitudinal study, demonstrates that dissolved oxygen (DO) concentrations in the Bronx River estuary remain consistently lower than those recorded from the Greenwich Cove estuary (Table 1). This is likely a result of ongoing urbanization proximal to the Bronx River estuary (Slattery, 2018). Consistent and long-lasting differences in dissolved oxygen levels from these two estuaries lend quantitative support to our claim that salt marsh grasses native to the Bronx River estuary are coping with challenges that rarely arise for their Greenwich Cove conspecifics.

**Table 1.** Estuarian water metrics. Average levels of dissolved oxygen (DO) over a 9-year survey, water temperature over a 7-year survey, and pH over a 5-year survey of the Bronx River estuary, NY and Greenwich Cove, CT. All water collections occurred in June and July, around low tide. Data are presented as mean ± SD. p < 0.0001 for dissolved oxygen concentrations, p=0.0063 for pH values.

| Site | Dissolved O<sub>2</sub> (mg/L) | Temperature (°C) | pH |
| --- | --- | --- | --- |
| Bronx River | $5.59 \pm .302$ | $21.93 \pm 1.05$ | $7.28 \pm 0.05$ |
| Greenwich Cove | $9.10 \pm .492$ | $24.62 \pm .807$ | $7.89 \pm .024$ |

Ultimately, the health of urban wetlands will depend on phenotypic adaptation of native species to the pressures introduced by anthropogenic processes (Alberti, 2015). Results of this study provide insight into the molecular mechanisms deployed by a dominant and foundational ecosystem provider that is facing consistent and long-term environmental challenge. Our results will help to inform future conservation and management decisions regarding critical urban plant species that must persist regardless of increasing urbanization and environmental degradation.

## 2. Materials and Methods

### 2.1 Water Collection and analyses

Concentrations of dissolved oxygen (DO) were obtained using the Winkler titration method. Oxygen was fixed on site using manganous sulfate, alkaline potassium iodide azide, and sulfamic acid. Sodium thio-sulfate was used to titrate the water sample with starch indicator to a clear endpoint.

### 2.2 Collection of plant material and heat shock

*Sporobolus alterniflorus* (formerly *Spartina alterniflora*) plants were removed from their native environments in Harding Lagoon near Soundview Park, Bronx River estuary, NY (40°48’35.6”N, 73°34’18.0”W), and at Todd’s Point estuary in Greenwich Cove, CT (41°0’ 30.8088’’N, 73°34’14.1709”W). Plants were taken along with their roots, soil, and native water at low tide. Plants were transported to the laboratory, where each plant was transplanted into native soil in a separate plastic cup with holes punched in the sides and base to allow for adequate aeration. All plants were equilibrated at approximately 23°C in a large basin of native water with aeration, for at least 24 hours. For the initial heat shock, four individual plants from each were incubated in a stand-up incubator at 42 ± 2°C for 30 minutes in pre-heated native water, and the incubator was left dark. Water was not aerated during treatment. Concurrently, four individual plants from each site (8 total plants), were incubated in their respective water at room temperature (21 ± 1°C) for 30 minutes in the dark without aeration to serve as controls. After treatments, all plants were moved to room temperature (both air and native water), ambient light, with no aeration to rest for 30 minutes. After resting, plants underwent a second treatment, heat shock or control, as described above. Post second treatment, all plants were moved to room temperature air and native water and allowed to rest overnight in ambient light with aeration. Grass samples were collected over multiple years, at the same time of year, and on the same tides, and as such represent appropriate biological replicates.

#### 2.3 DNA extraction

DNA was isolated using the Qiagen DNeasy Plant Mini Kit (Qiagen, #69104). Leaves were cut with clean dissecting scissors from near the stem of each individual plant and wiped clean of any dirt or debris. Plant matter (0.5 g) was taken for each treatment from each of four plants. Plant matter was pooled and cut into small pieces with clean dissecting scissors and crushed with a mortar and pestle and 400 μL AP1 (lysis buffer) + and 4 μL RNase. All contents were transferred to a micro centrifuge tube. Pipetting up and down further agitated samples. Samples were vortexed and incubated in a heat block at 65°C for 10 minutes, vortexing 2-3 times during incubation. The remaining steps were performed according to the Qiagen DNeasy Plant Mini Kit. DNA concentrations and quality were assessed using Nano Drop Microvolume Spectrophotometer (ThermoFisher Scientific). DNA samples were stored at -20°C in AE buffer from Qiagen DNeasy Plant Mini Kit.

### 2.4 Global DNA methylation analysis

Global genomic methylation levels were determined using 100 ng of DNA for each sample using the MethylFlash Methylated DNA Quantification Kit Colorimetric (Epigentek, #P-1034). The absolute methylation level was determined using a standard curve. The percentage of globally methylated DNA was calculated using absolute quantification from the Epigentek Protocol.

### 2.5 Protein extraction and concentration determination

Leaves were cut with clean dissecting scissors from near the stem of each individual plant and wiped clean of any dirt or debris. Plant tissue (500 mg) was pooled from four individuals from each site/condition. Tissue was immediately placed into lysis bags provided in the P-PER Plant Protein Extraction Kit (Thermo Scientific, #89803, Rockford, IL) and lysed according to provided instructions with the following modification: the volume of working solution was reduced to half of the recommended volume per sample. Protein solutions were stored at -20 °C for no longer than 4 weeks. Concentrations of proteins in plant leaves were determined according to the Pierce BCA Protein Assay Kit-Reducing Agent Compatible (Thermo Scientific, #23250, Rockford, IL).

### 2.6 Quantification of heat shock protein

Specific heat shock protein (HSP70) protein levels were quantified by Western blot (SDS-PAGE) analysis. Plant leaf protein samples (30-100 µg) were run on a 10% Mini-PROTEAN^®^ TGX™ Precast Protein Gel (Bio-Rad) and transferred to PVDF membranes, Ponceau stained to visualize protein transfer and evenly loaded across samples. Membranes were blocked for 1 hour at room temperature overnight at 4 °C in Superblock (Thermo-scientific, #37535, Rockford, IL). After blocking, blots were incubated in mouse HSP70 monoclonal antibody (1:1000, Enzo, #AZI-SPA-820, Plymouth Meeting, PA). After washing 3x 10min in TBS + Tween20, membranes were incubated in goat anti-mouse secondary antibody (1:10,000, Thermo-scientific, #31430, Rockford, IL) for 1 hour at room temperature. Membranes were washed 3 x 10min in TBS + Tween 20 before proteins were detected by Immuno-Star luminol-peroxide (Bio-Rad, #170-5070). Bio-Rad Chemi-doc was used to visualize and quantify proteins with densitometry.

### 2.7 DNA barcoding

DNA isolated from 12 organisms from each site (described above) was used to PCR amplify the universal plant gene *rbcL* using the following primers:

*rbcL*aF 5’-TGTAAAACGACGGCCAGTATGTCACCACAAACAGAGACTAAAGC-3’

*rbcL*a rev 5’CAGGAAACAGCTATGACGTAAAATCAAGTCCACCRCG-3’

Primers were a generous gift from the DNA Learning Center, Harlem, NY. PCR reactions were performed using GE illustra PURETaq Ready-To-Go PCR beads, 25μL of PCR reaction, 2μL of template DNA, and 11.5μL of primers. Reactions were run for 50 cycles: 30 seconds denaturation at 94°C, 45 seconds annealing at 54°C, and 45 seconds extending 72°C using Techne Genius Thermal Cycler. PCR amplicons were visualized on a 2% agarose gel using pBR322/BstNI molecular weight standards to ensure that products were the predicted size. PCR products were sent to Genewiz Inc. for sequencing. Sequences were aligned using Nucleotide BLAST. Percent similarity was ascertained using the CLUSTAL W (Thompson et al., 1994; Ni et al., 2012).

### 2.8 Statistical analysis

All water analysis data are presented as mean ± SD. p < 0.0001 for dissolved oxygen concentrations, p=0.0063 for pH values. Error bars and error ranges display ± S.E.M. as in Table 1 and Figure 2 respectively. P-values were determined through unpaired t-tests for statistical comparisons of two discrete populations as in Table 1. ANOVA tests for statistical comparisons of more than two experimental groups and/or discrete populations were performed as in Figure 2. P-values less than or equal to 0.05 are assumed to display statistical significance.

## 3. Results

### 3.1 Water analyses

Over a 9-year survey of the waters in the Bronx River estuary versus its less impacted counterpart in Greenwich Cove estuary, we collected physicochemical data regarding pH, temperature and dissolved oxygen levels. As shown in Table 1, the temperature and pH of the two sites are comparable. Temperature and pH for Bronx River estuary are 21.93 ±1.05 °C and 7.28 ± 0.05, while Greenwich Cove estuary are 24.62 ± 0.807 °C and 7.89 ± 0.024. However, dissolved oxygen concentrations at the two sites consistently vary widely, with the Bronx River estuary at 5.89 ± 0.30 mg/L and Greenwich Cove estuary at 9.69±2.2 mg/L. This difference is statistically significant (p < 0.0001). The National Oceanic and Atmospheric Administration (NOAA) National Estuarine Eutrophication Survey classifies water quality falling between 2 and 5 mg/L dissolved oxygen as stressed (Bricker et al., 1999). These data indicate that organisms residing in the Bronx River estuary are exposed to chronic, low oxygen levels, a condition that induces stress and metabolic dysfunction (Halem, et al., 2014; Ravaschiere et al., 2017).

### 3.2 Global methylation in response to stress for grasses from the differentially impacts sites

As shown in Figure 2, acute heat stress, as monitored by heat shock protein levels, results in inverse global DNA methylation profiles when comparing grasses collected from sites with distinctly different levels of environmental challenge. Percent global DNA methylation is increased by heat shock for grasses from the Bronx River estuary but is decreased by heat shock for Greenwich Cove organisms. These results are statistically significant (p values < 0.05). Interestingly, Bronx River grasses consistently demonstrate lower baseline global methylation levels as compared with their Greenwich Cove conspecifics (in Figure 1, compare Bronx and Greenwich controls).

**Figure 1:**
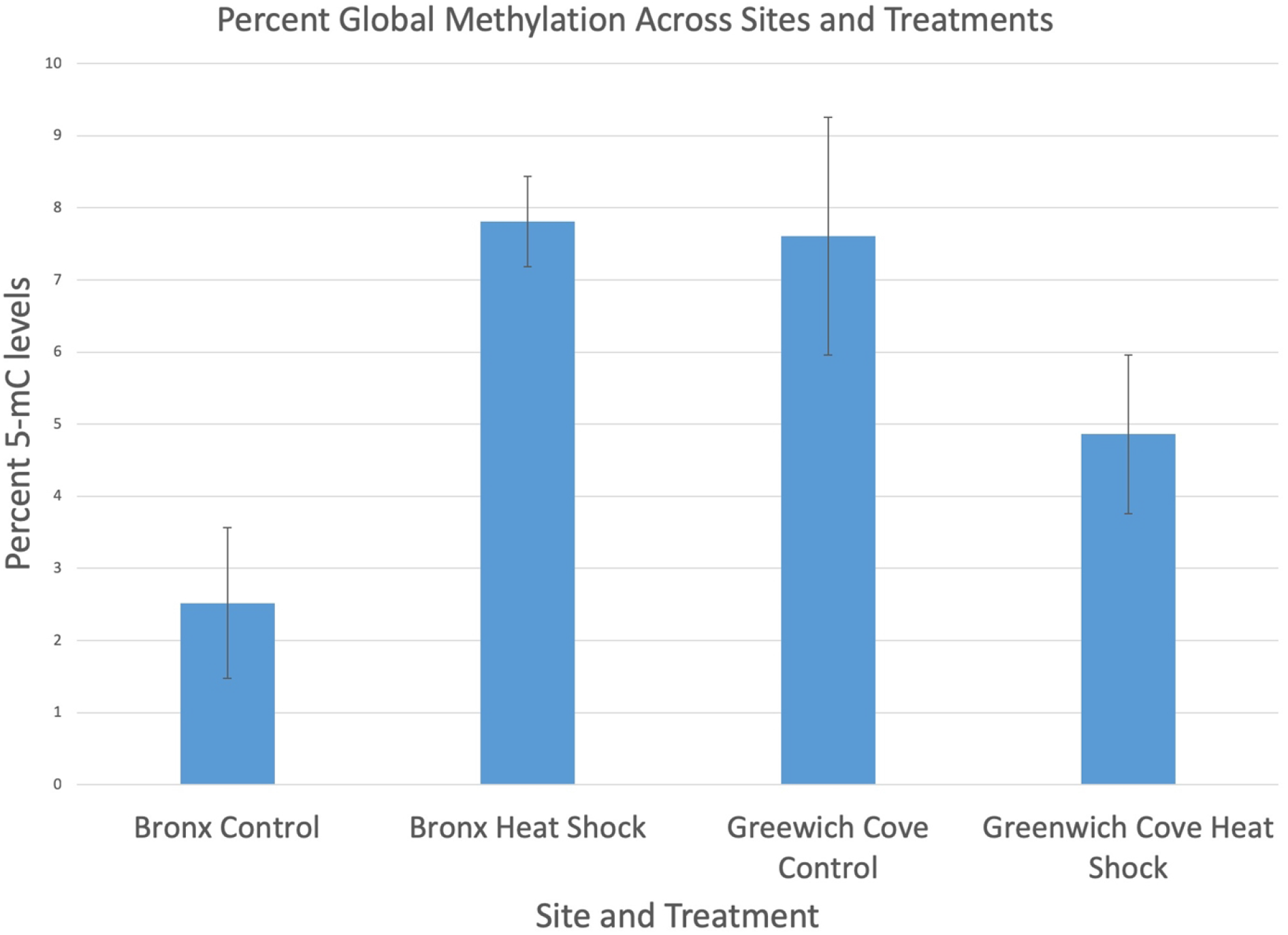
Quantification of global DNA methylation over two years. Average percent 5-methlycytosine (5-mC) of *S. alterniflorus* from the Bronx River and Greenwich Cove estuaries (n=4 for each group). The two populations launch inverse methylation responses in acute heat stress. Bronx individuals increase global methylation (p < 0.05), while Greenwich Cove individuals decrease global methylation, (p < 0.05). Statistical results are based on ANOVA tests.

**Figure 2:**
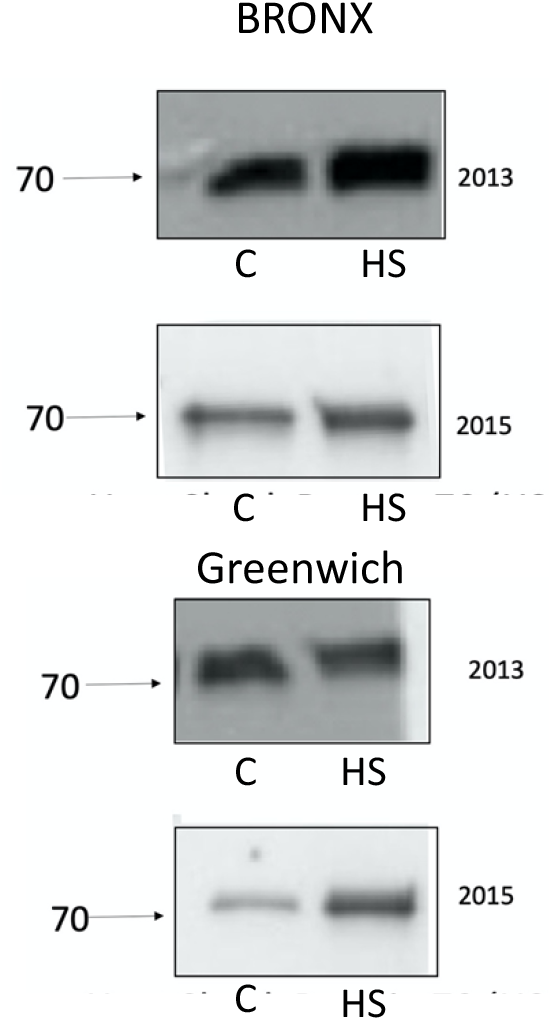
HSP70 Western blot analyses, *S. alterniflorus* (2013 and 2015), collected from Bronx River Estuary (top) and Greenwich Cove (bottom). Arrows point to 70 kDa HSP70 immunoreactive band. C= control and HS = heat shocked.

### 3.3 HSP70 protein levels indicate acute stress

Western blots analyses of heat shock immuno-reactive proteins bands (HSP70) across two years (Figure 2) show that heat shock proteins were successfully elicited in controlled laboratory settings in grasses from both sites.

### 3.4 Barcoding for species identification

Among *rbcL* chloroplast DNA gene sequences derived from individuals native to both the Bronx River and Greenwich Cove, CLUSTAL W alignment revealed negligible heterogeneity (0 -1%). In addition, CLUSTAL W alignment also showed insignificant (0 -1%) divergence in *rbcL* sequence within both populations. These results indicate that all specimens used in this study from both sites are members of the same species, either *Sporobolus alterniflorus* or *Sporobolus maritimus*. Based on geographical data *Sporobolus maritimus* is not native to the Bronx or Connecticut (USDA, plants database). Thus, we determined that *Sporobolus alterniflorus* is the species we analyzed at both sites, Greenwich Cove and Bronx River estuaries.

## 4. Discussion

Rapid global urbanization and its impact on the environment are fundamentally changing the course of evolution in organisms that represent our urban co-inhabitants (Hendry and Kinnison, 1999; Alberti, 2015; Donihue and Lambert, 2015; Johnson and Munshi-South, 2017). Plant life is critical to sustainable development of urban ecosystems (Barbier et al., 2011; Mexia et al., 2018). Thus, an important challenge in ecology is to gain a fuller understanding of the mechanisms by which plants cope with current unprecedented rates of environmental change. Information presented here contributes to a growing field of research that documents key molecular mechanisms by which plants adapt to the challenges of urbanization.

Studies that focus specifically on wild populations native to urban landscapes are especially well suited for predicting and conserving life in urban ecosystems. To date, few studies employ *in vivo* techniques to analyze the mechanisms of stress tolerance in wild populations as they respond to urbanization in their native habitats (Thiebau et al., 2019). For this study, two distinct wild populations of estuarine grasses were chosen as the biological models. *Sporobolus alterniflorus* was collected from two sites: the Bronx River, NY and Greenwich Cove, CT. These estuaries are located along an urban to suburban gradient, that features differential degrees of urbanization both quantitatively and historically (Rachlin et al., 2007; Halem et al., 2014; Ravaschiere et al., 2017). Physio-chemical analyses of estuary water, presented in Table 1, demonstrate that the Bronx River estuary consistently experiences long term, comparatively low dissolved oxygen, and these results are statistically significant. Hypoxic conditions in the Bronx River are associated with chronic stress response and metabolic disruptions in a foundation molluscan species (Halem, et al., 2014; Ravaschiere et al., 2017). Low dissolved oxygen primarily results from phosphorus and nitrogen run-off associated with wastewater and urban run-off. In addition, gas exchange in *Sporobolus alterniflorus* relies on direct access to dissolved oxygen from the aquatic environment via stems and roots that are often tidally submerged. These submerged parts of the plant, exposed to chronically low dissolved oxygen, must fulfill respiratory needs for the entire plant (Teal and Kanwisher, 1966). Thus, a low dissolved oxygen concentration directly inhibits vital metabolic processes and serves as a predictable indicator for urbanization and contamination.

Using estuarine grasses, from two distinct populations as described above, we simulated an abiotic stressor (heat stress) that is commonly associated with urbanization and climate change (Schlesinger et al., 2008; Fossog et al., 2013; Simon et al., 2016; Duarte et al., 2017). We did this by experimentally delivering a controlled heat shock, and monitoring stress response via immunochemical detection of the reliable stress indicator, heat shock protein 70 (HSP70). After heat shock was applied and induction of heat shock protein was confirmed by Western Blot analyses, we quantified changing levels of % 5-mC DNA methylation. This allowed us to monitor epigenetic modification as a response to acute stress in two differentially impacted plant populations. Figure 1 demonstrates that the Bronx River and Greenwich Cove grasses exhibited statistically different inverse DNA methylation profiles in response to the controlled application of acute heat stress. Western Blot analyses (Figure 2) confirm heat stress as indicated by the predicted and significant increases in heat shock protein (molecular weight 70 kD), and this response was documented across two years.

Results of our global DNA methylation analyses suggest that the two distinct populations of grasses counter the same heat stress by eliciting DNA methylation profiles that differ fundamentally and in biologically relevant ways. Our results show that grasses from the Bronx River estuary counter an acute stress via DNA hyper-methylation. Further studies are necessary to confirm our speculation that hyper-methylation may be an essential metabolic trade-off that globally shuts down gene expression, sparing only the basic housekeeping functions that are necessary for survival during acute stress. Whereas, for grasses native to the less impacted Greenwich Cove, application of acute stress elicits a reverse pattern, a hypo-methylation of the genome that may facilitate expression of those loci required for a successful acute stress response. In support of our findings, multiple recent studies demonstrate that individual plant populations counter similar environmental stressors in different ways, and this may be controlled at the epigenetic level (Saez-Laguna et al., 2014; Thiebaut et al., 2019; Rehman and Tanti, 2020).

The inverse epigenetic profiling that we report here between Bronx and Greenwich Cove grasses do not result from speciation. Our barcoding results demonstrate that grasses collected from these two geographically distinct populations have negligible differences in chloroplast *rbcL* gene sequence (Kress and Erikson, 2007). Very low rate of sequence divergence (1% or less) between individuals from the Bronx River and Greenwich Cove estuaries indicates that these individuals belong to the same species and are thus genetically identical. Differential DNA methylation within a species is now recognized as a critical indicator of short-term ecological experience (Rey et al., 2020). This notion of “ecological populations” will become essential in conservation biology with the recognition that epigenetics offers a key link between environmental change and phenotypic plasticity in wild populations (Rey et al., 2020).

Enhanced tolerance to stress could provide an evolutionary advantage for Bronx River grasses when confronted with the unpredictable challenges inherent to their native habitat. A phenomenon called defense priming conditions plants to better tolerate abiotic stressors, and this constitutes an important evolutionary benefit (Matinez-Medina et al., 2011; Crisp et al., 2016). This switch-like and reversible process offers sufficient genomic flexibility for sessile organisms, like plants, to respond rapidly to fluctuating environmental challenges (Meyer, 2008; Boyko and Kovalchuk, 2011; Arikan et al., 2018). Evidence suggests that defense priming improves fitness (Conrath et al., 2006; Matinez-Medina et al., 2011), and is regulated by epigenetic modification (Luna et al., 2012). In defense priming, a “stimulus” targets individual loci such that their methylation states become altered. This affects future accessibility of transcriptional machinery and gene expression eliciting a rapid and sustained response upon a later “triggering stimulus”. We hypothesize that long term hypoxia, as well as potential additional environmental challenges at the urbanized Bronx River estuary, constitute the priming “stimulus” for this particular population of grasses. Chromatin restructuring, specifically changes to DNA methylation patterns, is a mechanism by which plants engage in defense priming (Conrath et al., 2015; Savvides et al., 2016; Pastor et al., 2013), and here we demonstrate that two distinct plant populations deploy inverse patterns of global DNA methylation when encountering the same acute stressor. These results suggest that grasses from the Bronx River Estuary may be engaged in defense priming. Our research contributes to rapidly growing body of evidence demonstrating that successful strategies for coping with rapid environmental change do not rely solely on random heritable changes in DNA sequence. For sessile species such as plants, critical evolutionary adaptations are propelled by epigenetic change (Youngson et al., 2008; Holeski et al., 2012; Robertson et al., 2017; Thiebau et al., 2019; Whittle et al., 2019; Rey el al., 2020).

In conclusion, current predictions regarding species tolerance to urbanization and climate change are based on the assumption that all members of a species elicit the same, or similar responses to an environmental stressor (Sih et al., 2011). However, data presented here corroborate growing evidence suggesting that populations separated geographically or by differential stressors indeed demonstrate key distinct coping mechanisms (Fossog et al., 2013; Halem et al., 2014; Ravaschiere et al., 2017; De Almeida Duarte et al., 2017). In other words, members of the same species may utilize diverging mechanisms in order to continue to thrive in the face of rapid and challenging environmental stressors. In addition, results presented here suggest that stress priming by epigenetic modification may enhance fitness in grasses native to the heavily impacted Bronx River estuary. These complex molecular mechanisms of biological adaptation deserve our attention so that we may better understand the ways in which particular populations develop unique adaptive responses that are specifically suited to unique ecological challenges.

As molecular ecologists and evolutionary biologists, we must accept the notion that environmentally induced adaptation is a key feature to sustaining resilience and biodiversity in rapidly expanding urban areas. As conservationists and urban planners, we must gain better understandings of the unique epigenetic mechanisms by which foundational, ecological service providers adapt and evolve to tolerate the pressures of urbanization. Future studies are required in order to identify genes that are differentially methylated in response to stress in *Sporobolus alterniflorus*, a species that is vital to urban sustainability. These studies await the full sequence analysis and annotation of the *Sporobolus alterniflorus* genome. In addition, future studies should be initiated to more fully understand the intricacies of adaptation in populations that remain chronically exposed to low oxygen and other stressors associated with urbanization.

## Acknowledgements

We thank Riverdale Country School, Dominic Randolph and Kelley Nicholson-Flynn for continued support of the Lisman Laboratories. We acknowledge Zachery Halem for his help with planning discussions and analyses. We are grateful to Julia Gardner who assisted with literature searches, and to Travis Brady, Emma Dietz, Juliette Egan, Jacob Katz, Eli Sands, Shae Simpson, and Knut Vanderbush for their invaluable assistance in the laboratory. We thank Melissa Lee and the Harlem DNA Center for donating PCR primers and advice. We thank Dan Luciano who generously shared equipment and expertise.

